# A new and highly effective method for predicting T-cell response targets implemented on SARS-CoV-2 data

**DOI:** 10.1101/2022.09.09.507307

**Authors:** Jaroslav Flegr, Daniel Zahradník

## Abstract

Computational T-cell epitope prediction is essential in many immunological projects, including the development of vaccines. T-cells of immunocompetent vertebrate hosts can recognize as non-self only peptides which are present in the parasite’s proteins and absent in the host’s proteins. This basic principle allows us to predict which peptides can elicit T-cells’ response. We built on the fact that the specificity of T-cells reacting to SARS-CoV-2 antigens has been recently mapped in detail. Using Monte Carlo tests, we found that empirically confirmed peptides that stimulate T-cells contain an increased fraction of pentapeptides, hexapeptides, and heptapeptides which are not found in the human proteome (p < 0.0001). Similarly, hexapeptides absent in human proteins were overrepresented in peptides that elicited T-cell response in a published empirical study (p = 0.027). The new theory-based method predicted T-cell immunogenicity of SARS-CoV-2 peptides four times more effectively than current empirically based methods.

## Introduction

The adaptive immune system of vertebrates discriminates between self and non-self antigens primarily by detecting peptides originating from non-self proteins. The detection of unknown molecules (or any previously unknown entities) might seem a challenging or even unsolvable problem, especially in comparison with the detection of known molecules or entities, but evolution found an elegant way of solving this principally tricky task. It is based on an interplay of T cells, MHC proteins, and antigen-presenting cells (1).

Nearly all vertebrate nucleated cells continuously digest a sample of all their endogenous proteins and present on their surface for inspection by T cells a subsample of the resulting peptides, that is, the peptides which have an affinity to some variant of their MHC class I proteins. Similarly, professional antigen-presenting cells, such as dendritic cells or B-cells, digest a sample of all proteins which were transported to their interior by phagocytosis or endocytosis, and present on their surface a subsample of this sample, that is, peptides with affinity to some variant of their MHC class II proteins. Antigen-presenting cells cannot discriminate between self and non-self peptides: both types of peptides are therefore presented on their surface for inspection by T cells. Clones of T-cells, each recognizing a different peptide – or a small group of peptides – kept by non-covalent bonds in the groove of MHC proteins, likewise cannot discriminate between self and non-self peptides. But during maturation, they pass through the thymus, where T cells with a strong enough affinity to any peptide attached to the MHC molecules (peptides originated from self-proteins) either die or differentiate into a specialized type of T cells – regulatory T cells (2–4). Due to this combination of negative selection and elimination of T cells which bear receptors with insufficient affinity to any MHC-peptide complex, most mature cytotoxic and helper T cells patrolling in our bodies recognize only non-self peptides, that is, peptides which are not presented in the thymus.

In their coevolutionary arms race with hosts, parasitic organisms gradually eliminate all unnecessary pentapeptides from their peptide vocabulary. It has been demonstrated that a parasitic lifestyle affects the size of organisms’ pentapeptide vocabulary, i.e., the number of different pentapeptides in their proteins, more strongly than the size or complexity of their proteome (5). To avoid recognition by host’s T cells, parasites with a broad host specificity eliminate all unnecessary pentapeptides, that is, they reduce the size of their peptide vocabularies. Parasites characterized by a narrow host specificity substitute peptides not present in the proteins of their natural host species with those which are present there. In particular, they mutate these peptides, potential targets of T-cell recognition, into peptides present in the peptide vocabulary of their host species. In viruses, bacteria, and parasitic protozoa, this evolutionary process can take place both within the entire metapopulation or in particular infrapopulations, i.e., in the populations of parasites within individual hosts. The process can be therefore relatively rapid and can even affect the progress of a disease in an individual patient. The resulting peptide vocabulary mimicry could be partly responsible for the phenomenon of host specificity, i.e., for the fact that most parasite species have a limited range of potential host species (6).

Similarity between pentapeptide and hexapeptide vocabularies can reveal the species of the natural host of a particular parasite species. This kind of analysis had recently shown that the natural host of the ancestor of SARS-CoV-2 was probably a horseshoe bat, while the natural host of donor of its spike gene was human (7). The same study also showed that the donor of most protein-coding genes of SARS-CoV-2 were probably parasitized treeshrews in recent past, while the donor of the spike gene had probably recently parasitized rats.

In this study, we tested a critical prediction of the peptide vocabulary mimicry theory: we investigated whether the immunogenicity of peptides of viral origin can be predicted based on the content of pentapeptides and hexapeptides which are not present in their host’s peptide vocabularies. We took advantage of a recently published list of 734 peptides which elicited a specific immune response of cytotoxic or helper T cells isolated from 99 post-Covid patients (8). Using *in silico* methods, we sought the intersecting points between the list of real (empirically identified) T-cell response targets and a list of potential T-cell recognition targets, that is, peptides present in SARS-CoV-2 but absent in the human proteome.

## Results

### 3.1. How many potential targets of human T-cell recognition are there in SARS-CoV-2 proteins?

We prepared penta-, hexa-, and heptapeptide vocabularies for humans and SARS-CoV-2 as well as lists of peptides present in the SARS-CoV-2 proteome but absent in the human proteome, that is, lists of potential targets of T-cell recognition. The genetic distance between SARS-CoV-2 and humans is considerable, which is also why most viral peptides longer than six amino acids were absent in human proteins. Our analyses found that 983 (10.2%) of all 9,609 pentapeptides, 7,091 (73.7%) of all 9,620 hexapeptides, and 9,334 (97.2%) of all 9,598 heptapeptides of SARS-CoV-2 were absent in human peptides and therefore could serve as potential targets of human T-cell recognition (Fig. 1). The density of potential pentapeptide targets of T-cell recognition was lower in the spike, which originated from the human-adapted SARS-CoV-2 ancestor (7), than in ten other SARS-CoV-2 proteins, which originated from a horseshoe bat-adapted ancestor (8.75% vs. 11.07%, Chi^2^ = 5.80, p = 0.016) (7). No difference in the density of potential targets was observed for hexapeptides (7.43% vs. 7.44%, Chi^2^ = 0.01, p = 0.922) or heptapeptides (9.79% vs. 9.72%, Chi^2^ = 1.84, p = 0.175).

**Fig. 1.**
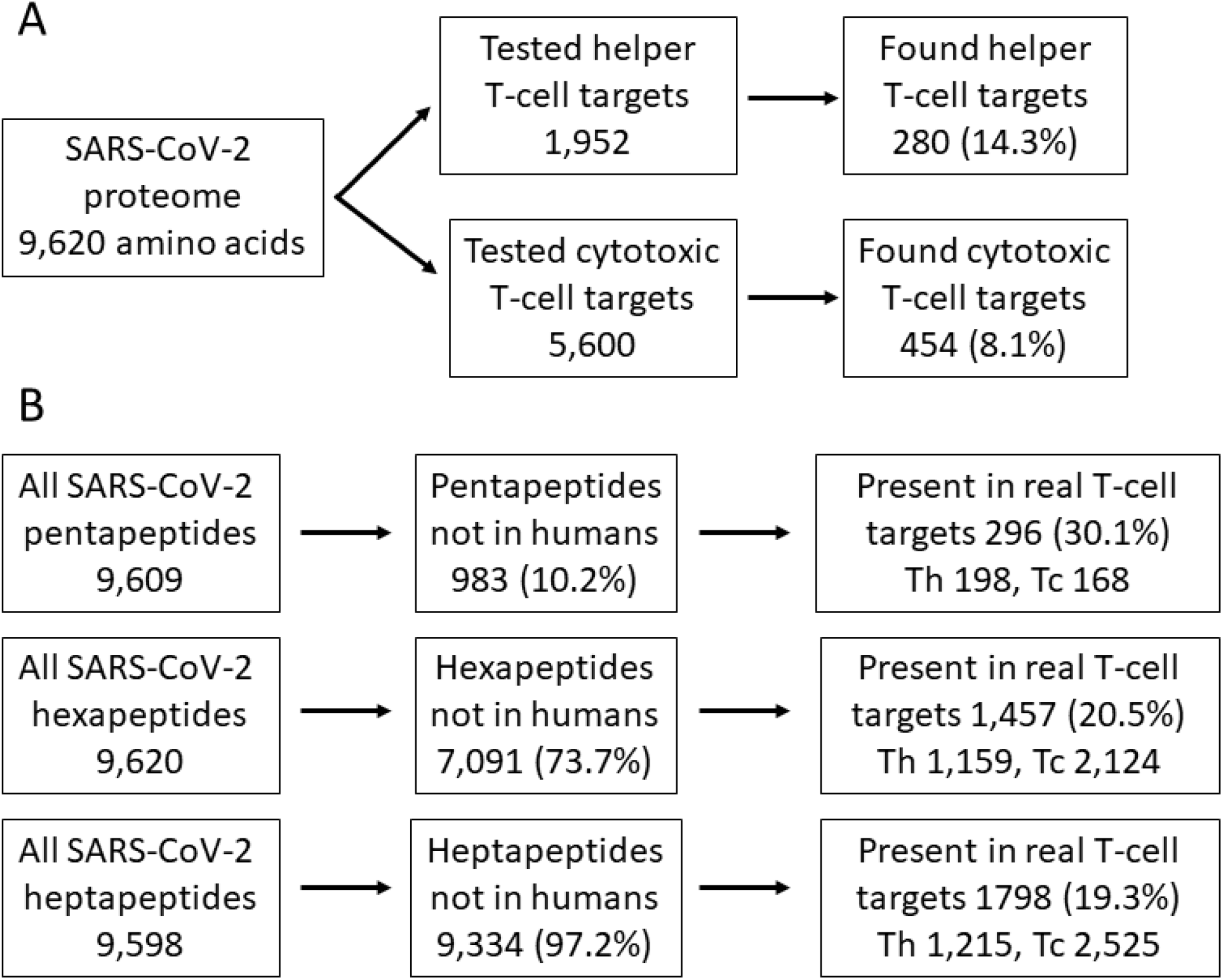
Classification and representation of targets of T-cell response and potential targets of T-cell recognition Part A offers a basic outline of genuine targets of helper and cytotoxic T-cell response in empirically tested SARS-CoV-2 peptides (8). Part B shows a basic outline of potential targets of T-cell recognition contained in peptides eliciting T-cell response in all potential targets of T-cell recognition and in all SARS-CoV-2 peptides.

### 3.2. What fraction of empirically found immunogenic peptides contained potential targets of T-cell immune recognition?

We calculated the fraction of peptides identified by Tarke *et al*. as real targets of T-cell response which contained at least one of our potential penta-, hexa-, and heptapeptide targets of T-cell recognition. Then we used a Monte Carlo test to estimate whether this fraction is larger than the equivalent fraction calculated for a case where potential targets of T-cell recognition are selected from SARS-CoV-2 peptide vocabularies randomly (or, alternatively, where our method of recognition of T-cell targets does not work).

Tarke *et al*. tested 1,952 15mer peptides covering whole SARS-CoV-2 proteomes and found 280 (14.3%) peptides which were, according to their criteria, the targets of actual helper T-cell response (Fig. 2A). Using predictive algorithms, they found 5,600 potential targets of response by cytotoxic T cells; in subsequent empirical essays, they found that 454 (8.11%) of them actually elicited a response by cytotoxic T cells. We found that 293 (39.9%) of all 734 targets of response by helper or cytotoxic T cells contained some of our 983 potential pentapeptides, 727 (99.0%) some of our 7,091 potential hexapeptides, and 734 (100%) some of our 9,334 heptapeptide targets of T-cell recognition. Monte Carlo tests showed that the potential pentapeptide and hexapeptide T-cell targets were strongly overrepresented in peptides which were recognized by helper T cells. When 983 pentapeptides and 7,091 hexapeptides were selected 10,000 times randomly from all SARS-CoV-2 pentapeptides and hexapeptides, only on average 75.18 (10.2%) and 541.07 (73.7%) peptides eliciting T-cell response contained some of the randomly selected pentapeptides and hexapeptides, respectively. The difference in representation of our potential targets of T-cell recognition and the ‘pseudotargets’ of randomly selected sets in peptides that really elicited T-cell response was highly significant for both pentapeptides (39.9% vs. 10.2%, p < 0.0001, Cohen’s d = 27.50) and hexapeptides (99.0% vs. 73.7%, p < 0.0001, Cohen’s d = 16.2). In fact, pseudotargets of all of the 10,000 random sets of peptides were represented less frequently in peptides eliciting T-cell response than in the genuine set of our potential targets of T-cell recognition. Potential heptapeptide targets of recognition were present in all peptides eliciting a T-cell response, while randomly selected pseudotargets were present only in 97.0% of such peptides (Fig. 2). Again, the overrepresentation of genuine potential targets in peptides eliciting a T-cell response was highly significant (100% vs. 97.0%, p < 0.0001, Cohen’s d = 4.98). The effect size of all observed effects was very large: according to the widely used Cohen’s nomenclature, all effects with Cohen’s d > 0.8 are classified as large (9).

**Figure 2.**
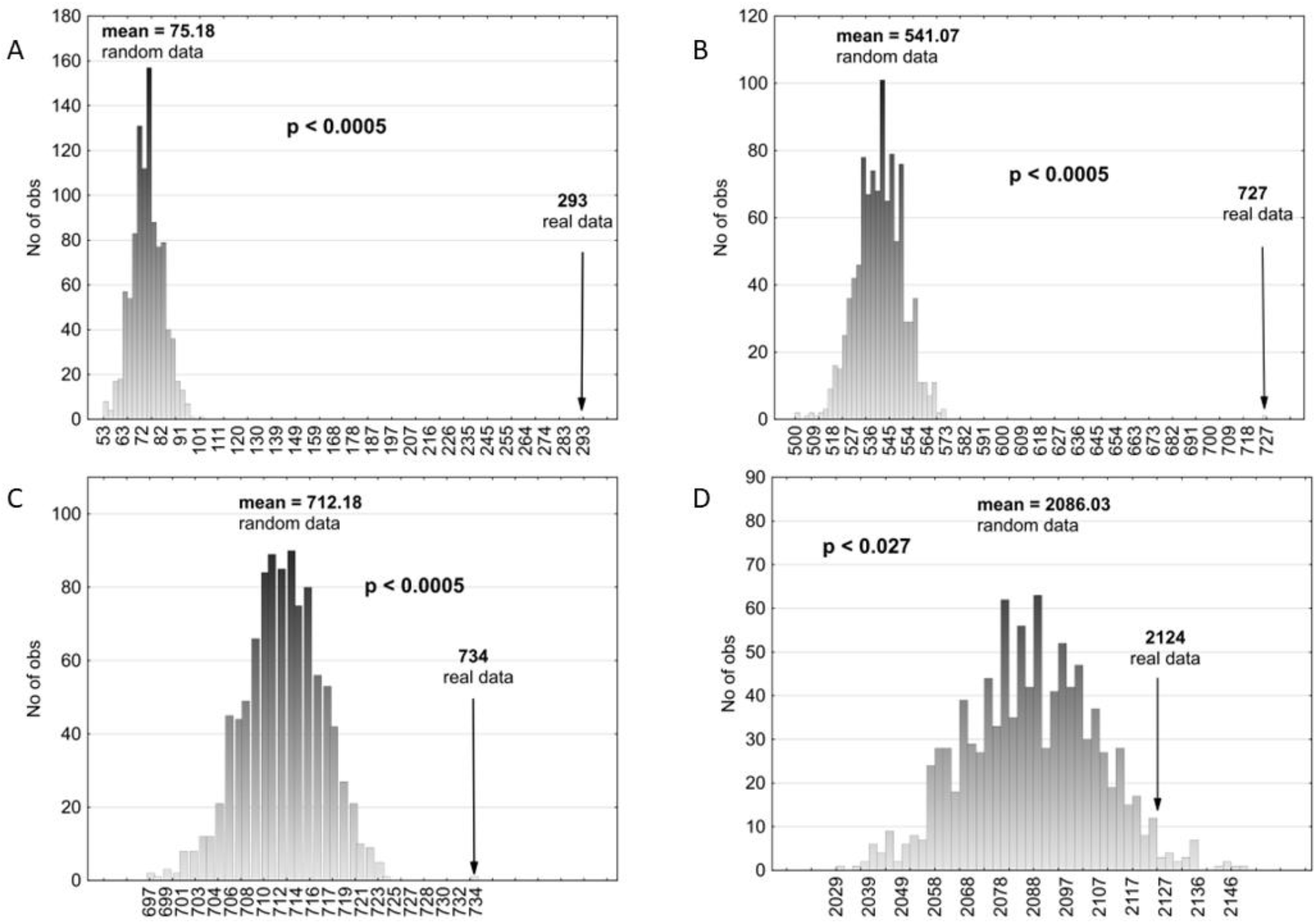
Results of the Monte Carlo test Parts A, B, and C show an overrepresentation of potential targets which were present in peptides eliciting a T-cell response within all potential targets of T-cell recognition (penta-, hexa-, and heptapeptides, respectively). Part D shows an overrepresentation of peptides eliciting a T-cell response which contained potential targets T-cell recognition (hexapeptides).

Post hoc analyses were performed separately for targets of response by helper and cytotoxic T cells which originated from the spike protein and the rest of the SARS-CoV-2 proteome (see Table 1). Analyses showed that the effect – that is, the overrepresentation of pentapeptide and hexapeptide potential targets of recognition in real targets of T-cell response – was about twice stronger for helper T cells than for cytotoxic T-cell targets, and it was weaker for the spike protein than for other proteins. The frequency of immunogenic peptides containing potential targets of T-cell recognition was lower in the spike protein than in other SARS-CoV-2 proteins (see Table 1).

**Table 1.** Representation of potential targets of T-cell recognition in peptides that elicited a T-cell response This table shows the number of peptides which elicit a T-cell response (3^rd^ column), those which contain potential targets of T-cell recognition (4^th^ column), representation of these targets in peptides that elicit a T-cell response (5^th^ column), and the results of the Monte Carlo test. In particular, the last five columns show the mean number of peptides containing randomly selected pseudotargets, representation of these pseudotargets in peptides eliciting a T-cell response, standard deviation of the number of peptides containing randomly selected pseudotargets, significance (p) of the one-sided Monte Carlo test, and one-sample Cohen’s d showing the strength of the effect reflecting the difference between columns 4 and 6. Significance under 0.0001 is marked as 0.000.

|  |  | No.<br>targets | With<br>genuine<br>targets | With<br>genuine<br>targets % | With<br>random<br>targets | With<br>random<br>targets % | sd<br>random<br>targets | p | Cohen's<br>d |
| --- | --- | --- | --- | --- | --- | --- | --- | --- | --- |
| T cells,<br>whole virus | 5-peptides | 734 | 293 | 39.9% | 75.18 | 10.2% | 7.92 | 0.000 | 27.50 |
|  | 6-peptides | 734 | 727 | 99.0% | 541.07 | 73.7% | 11.48 | 0.000 | 16.20 |
|  | 7-peptides | 734 | 734 | 100.0% | 712.18 | 97.0% | 4.38 | 0.000 | 4.98 |
| Th cells,<br>whole virus | 5-peptides | 280 | 155 | 55.4% | 28.72 | 10.3% | 5.04 | 0.000 | 25.04 |
|  | 6-peptides | 280 | 280 | 100.0% | 206.41 | 73.7% | 7.30 | 0.000 | 10.10 |
|  | 7-peptides | 280 | 280 | 100.0% | 271.65 | 97.0% | 2.78 | 0.000 | 3.01 |
| Tc cells,<br>whole virus | 5-peptides | 454 | 138 | 30.4% | 46.50 | 10.2% | 6.31 | 0.000 | 14.50 |
|  | 6-peptides | 454 | 447 | 98.5% | 334.65 | 73.7% | 9.24 | 0.000 | 12.16 |
|  | 7-peptides | 454 | 454 | 100.0% | 440.45 | 97.0% | 3.53 | 0.000 | 3.84 |
| T cells,<br>spike<br>protein | 5-peptides | 247 | 86 | 34.8% | 21.63 | 8.8% | 3.96 | 0.000 | 16.26 |
|  | 6-peptides | 247 | 241 | 97.6% | 183.40 | 74.3% | 6.14 | 0.000 | 9.40 |
|  | 7-peptides | 247 | 247 | 100.0% | 241.76 | 97.9% | 2.04 | 0.002 | 2.58 |
| Th cells,<br>spike<br>protein | 5-peptides | 92 | 46 | 50.0% | 8.02 | 8.7% | 2.63 | 0.000 | 14.44 |
|  | 6-peptides | 92 | 92 | 100.0% | 68.36 | 74.3% | 4.05 | 0.000 | 5.84 |
|  | 7-peptides | 92 | 92 | 100.0% | 90.03 | 97.9% | 1.35 | 0.135 | 1.46 |
| Tc cells,<br>spike<br>protein | 5-peptides | 155 | 40 | 25.8% | 13.53 | 8.7% | 3.25 | 0.000 | 8.14 |
|  | 6-peptides | 155 | 149 | 96.1% | 115.16 | 74.3% | 5.13 | 0.000 | 6.60 |
|  | 7-peptides | 155 | 155 | 100.0% | 151.72 | 97.9% | 1.69 | 0.031 | 1.94 |
| T cells,<br>other<br>proteins | 5-peptides | 487 | 207 | 42.5% | 53.84 | 11.1% | 6.62 | 0.000 | 23.13 |
|  | 6-peptides | 487 | 486 | 99.8% | 362.47 | 74.4% | 9.20 | 0.000 | 13.43 |
|  | 7-peptides | 487 | 487 | 100.0% | 473.34 | 97.2% | 3.50 | 0.000 | 3.94 |
| Th cells,<br>other<br>proteins | 5-peptides | 188 | 109 | 58.0% | 20.83 | 11.1% | 4.22 | 0.000 | 20.90 |
|  | 6-peptides | 188 | 188 | 100.0% | 139.93 | 74.4% | 5.92 | 0.000 | 8.12 |
|  | 7-peptides | 188 | 188 | 100.0% | 182.73 | 97.2% | 2.25 | 0.004 | 2.34 |
| Tc cells,<br>other<br>proteins | 5-peptides | 299 | 98 | 32.8% | 33.90 | 11.3% | 5.21 | 0.000 | 12.45 |
|  | 6-peptides | 299 | 298 | 99.7% | 222.54 | 74.4% | 7.26 | 0.000 | 10.40 |
|  | 7-peptides | 299 | 298 | 99.7% | 290.58 | 97.2% | 2.74 | 0.000 | 3.07 |

### 3.3 What fraction of potential T-cell targets was present in the known targets of T-cell response?

Tarke *et al*. found 280 15mer peptides recognized by Th cell and 454 peptides 9–14 amino acids-long which elicited Tc-cell response (Fig. 2A). We found that 296 (30.1%) of the 983 identified potential pentapeptide targets, 2,124 (30.0%) of the 7,091 identified potential hexapeptide targets, and 2,525 (27.1%) of the 9,334 heptapeptide targets were present either among the empirically identified peptides stimulating the Th cells (pentapeptides: 198; hexapeptides: 1,457; heptapeptides: 1,798) or among Tc-cell epitopes (pentapeptides: 168, hexapeptides: 1,159, heptapeptides: 1,215), see Fig. 2B. Some of them, namely 70 pentapeptides, 492 hexapeptides, and 488 heptapeptides, were even present in both Th-cell and Tc-cell epitopes.

We used a Monte Carlo test to examine whether the potential penta-, hexa-, and heptapeptide targets were present among the genuine targets of T-cell response significantly more often than random sets of SARS-CoV-2 penta-, hexa-, and heptapeptides. We compared the representation of potential penta-, hexa-, and heptapeptide targets in the genuine targets of T-cell response with the representation of 983 pentapeptides, 7,091 hexapeptides, and 9,334 heptapeptides 10,000 times randomly selected from the SARS-CoV-2 proteome (Fig. 1). The results showed that our *in silico*-identified hexapeptides, but not pentapeptides or heptapeptides, were overrepresented in the empirically identified targets of T-cell recognition in post-Covid patients (hexapeptides: genuine peptides 2,124, random peptides 2,186.03, p = 0.027, Cohen’s d = 1.97; pentapeptides: genuine peptides 296, random peptides 309.95, p = 0.834, Cohen’s d = 1.02; heptapeptides: genuine peptides 2,525, random peptides 2,526.49, p = 0.658, Cohen’s d = 0.47) (Fig. 3D). Post hoc tests performed separately for targets of cytotoxic and helper T cells showed that the results were significant for hexapeptides recognized by cytotoxic T cells (p = 0.040, Cohen’s d = 1.77, genuine 1,159 vs. random 1,130.33) but not for those recognized by helper T cells (p = 0.075, Cohen’s d = 1,44, genuine 1,457 vs. random 1,433.07). The effects were weaker and less numerous than those described in section 3.2. but remained very large according to Cohen’s classification.

**Fig. 3.**
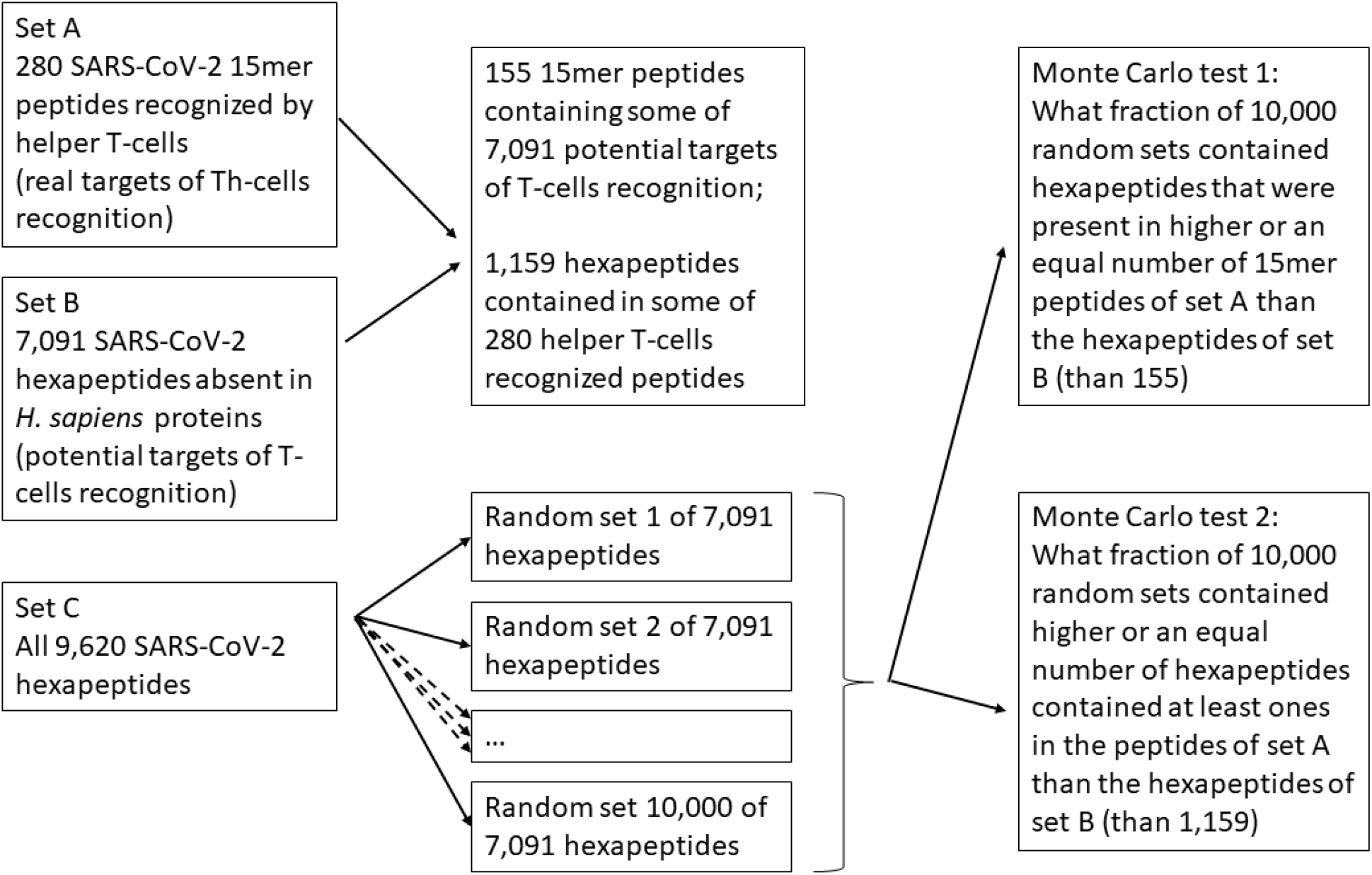
Schedule of Monte Carlo tests used in the present study This schedule uses hexapeptides as an example. The same approach was applied to pentapeptides and heptapeptides.

Table 2 shows the results of Monte Carlo tests performed separately for the whole virus (the upper section), for its spike protein originating from the human-adapted virus (the middle section), and for the rest of the virus, i.e., the ten genes which originated from the horseshoe bat-adapted SARS-CoV-2 progenitor (the bottom section). Overrepresentation of potential hexapeptide targets of T-cell recognition in peptides eliciting a T-cell response was significant only for those immunogenic peptides originating in the spike protein which elicited a helper T-cell response. Analyses of proteins originating from the other ten proteins showed an overrepresentation of potential heptapeptide targets of T-cell recognition which were present in some peptides stimulating a T-cell response.

**Table 2.** Representation of potential targets which were present in peptides eliciting T-cell response within all potential targets of T-cell recognition This table shows the number of potential targets of T-cell recognition (3^r^ column), the number of those that were really present in peptides eliciting T-cell response (4^th^ column), their representation among all potential targets of T-cell recognition (5^th^ column), and the results of Monte Carlo test. In particular, the last five columns show the mean number of randomly selected pseudotargets which were present in peptides eliciting a T-cell response, their representation within all pseudotargets, standard deviation of the number of pseudotargets present in peptides eliciting a T-cell response, significance of the one-sided Monte Carlo test, and a one-sample Cohen’s d showing the strength of the effect, thus reflecting the difference between columns 4 and 6.

|  |  | No.<br>targets | Within<br>genuine | Within<br>genuine % | Within<br>random | Within<br>random % | sd<br>random | p | Cohen's<br>d |
| --- | --- | --- | --- | --- | --- | --- | --- | --- | --- |
| T cells,<br>whole virus | 5-peptides | 983 | 296 | 30.1% | 309.95 | 31.5% | 13.71 | 0.834 | 1.02 |
|  | 6-peptides | 7091 | 2124 | 30.0% | 2086.03 | 29.4% | 19.29 | 0.027 | 1.97 |
|  | 7-peptides | 9334 | 2525 | 27.1% | 2526.49 | 27.1% | 7.39 | 0.658 | 0.47 |
| Th cells,<br>whole virus | 5-peptides | 983 | 198 | 20.1% | 206.35 | 21.0% | 11.98 | 0.743 | 0.70 |
|  | 6-peptides | 7091 | 1457 | 20.5% | 1433.07 | 20.2% | 16.64 | 0.075 | 1.44 |
|  | 7-peptides | 9334 | 1798 | 19.3% | 1802.75 | 19.3% | 6.54 | 0.751 | 0.73 |
| Tc cells,<br>whole virus | 5-peptides | 983 | 168 | 17.1% | 183.2 | 18.6% | 11.45 | 0.906 | 1.33 |
|  | 6-peptides | 7091 | 1159 | 16.3% | 1130.33 | 15.9% | 15.92 | 0.040 | 1.77 |
|  | 7-peptides | 9334 | 1215 | 13.0% | 1226.34 | 13.1% | 5.56 | 0.969 | 2.01 |
| T cells,<br>spike protein | 5-peptides | 111 | 78 | 70.3% | 80.12 | 72.2% | 4.51 | 0.646 | 0.47 |
|  | 6-peptides | 942 | 663 | 70.4% | 650.05 | 69.0% | 7.22 | 0.032 | 1.79 |
|  | 7-peptides | 1240 | 795 | 64.1% | 795.67 | 64.2% | 2.44 | 0.515 | 0.27 |
| Th cells,<br>spike protein | 5-peptides | 111 | 65 | 58.6% | 61.16 | 55.1% | 5.03 | 0.198 | 0.76 |
|  | 6-peptides | 942 | 518 | 55.0% | 501.46 | 53.2% | 7.83 | 0.015 | 2.11 |
|  | 7-peptides | 1240 | 621 | 50.1% | 618.52 | 49.9% | 2.54 | 0.118 | 0.98 |
| Tc cells,<br>spike protein | 5-peptides | 111 | 42 | 37.8% | 49.33 | 44.4% | 5.02 | 0.915 | 1.46 |
|  | 6-peptides | 942 | 369 | 39.2% | 364.09 | 38.7% | 7.58 | 0.238 | 0.65 |
|  | 7-peptides | 1240 | 391 | 31.5% | 393.41 | 31.7% | 2.35 | 0.793 | 1.03 |
| T cells,<br>other genes | 5-peptides | 614 | 218 | 35.5% | 234.77 | 38.2% | 11.35 | 0.924 | 1.48 |
|  | 6-peptides | 4132 | 1462 | 35.4% | 1455.64 | 35.2% | 15.51 | 0.336 | 0.41 |
|  | 7-peptides | 5387 | 1730 | 32.1% | 1721.19 | 32.0% | 5.76 | 0.049 | 1.53 |
| Th cells,<br>other genes | 5-peptides | 614 | 133 | 21.7% | 146.44 | 23.9% | 9.98 | 0.902 | 1.35 |
|  | 6-peptides | 4132 | 940 | 22.7% | 945.9 | 22.9% | 13.72 | 0.653 | 0.043 |
|  | 7-peptides | 5387 | 1177 | 21.8% | 1170.15 | 21.7% | 5.08 | 0.071 | 1.35 |
| Tc cells,<br>other genes | 5-peptides | 614 | 126 | 20.5% | 136.14 | 22.2% | 9.81 | 0.837 | 1.03 |
|  | 6-peptides | 4132 | 790 | 19.1% | 776.19 | 18.8% | 12.63 | 0.129 | 1.09 |
|  | 7-peptides | 5387 | 824 | 15.3% | 816.38 | 15.2% | 4.45 | 0.029 | 1.71 |

## Discussion

Our *in silico* method predicted a subpopulation of pentapeptides and hexapeptides with the potential to induce a T-cell response in humans. These peptides were overrepresented in peptides that had been experimentally identified as targets of T-cell response in post-Covid patients.

About 70% of potential targets of T-cell recognition were absent in all peptides which had been recognized as targets of T-cell response by immunological assays. Many of them were probably absent for trivial reasons, such as that the corresponding peptides were not tested in the Tarke *et al*. study or did not bind to any variant of MHC proteins of the 99 post-Covid patients involved in the study. There are, however, at least two other, less trivial explanations. To fit in the groove of MHC proteins and to be kept there firmly enough, specific amino acids must surround the peptide recognized by a T-cell. Moreover, a potential immune response target can become an actual immune response target only if it is present in the protein expressed intensively enough in the virus-infected cells. Tarke *et al*. showed that most peptides stimulating a T-cell response were present in just 4 out of 22 proteins, most frequently in the spike protein. Potential targets of T-cell recognition present in the weakly expressed or less immunogenic proteins therefore had a low chance of being detected in an empirical study.

Our study confirmed that a strikingly higher fraction of predicted targets (70%) was observed in empirically detected T-cell immunogenic peptides originating from the highly expressed and highly immunogenic spike protein than in the whole SARS-CoV-2 proteome (30%). This explanation thus seems to be the most probable of all the (non-exclusive) explanations mentioned just above. In this context, it is essential to bear in mind that the density of potential pentapeptide targets of T-cell recognition was lower in the highly expressed, highly immunogenic spike protein than in other SARS-CoV-2 proteins. This difference in densities is in agreement with earlier results, which suggest that SARS-CoV-2 is a chimera whose spike protein had originated from a donor which was initially adapted to a human host (and therefore lacks most peptides which are absent in the human peptidome), while the rest of the proteome originated from another species of coronavirus that was initially adapted to horseshoe bats (7). In typical parasites of nonchimeric origin, the densities of T-cell recognition targets will be similar in all parts of the proteome. Hence also the correlation between immunogenicity of an individual protein and the density of its potential T-cell recognition targets will probably be higher for such parasites than for the SARS-CoV-2.

All peptides that elicited T-cell immune response in post-Covid patients contained at least one predicted hexapeptide and heptapeptide target of immune recognition, and about one half of the immunogenic peptides contained at least one of the less numerous pentapeptide targets. Results of the Monte Carlo tests show that such overrepresentation is highly improbable and cannot happen by chance. In fact, peptides of none of the 10,000 randomly selected sets of peptides were similarly overrepresented in the 280 helper T-cell epitopes. This stronger overrepresentation of hexapeptides than pentapeptides (Table 2) might seem paradoxical. Intuitively, we would expect the presence potential hexapeptide target to be necessarily associated with the existence of a pentapeptide target. This principle, however, applies only in the reverse direction: the presence of potential pentapeptides is necessarily associated with the presence of hexapeptide targets. A comparison of our lists of potential T-cell recognition targets showed that most of the potential hexapeptide T-cell targets (76.7%) contained only pentapeptides which are present in the human proteome. As a result, a significant fraction of potential hexapeptide targets contains no potential pentapeptide target.

Analyses of overrepresentation of peptides containing at least one potential target of T-cell recognition in peptides that elicited T-cells response within all peptides that elicited T-cell response, as well as the lower density of potential pentapeptide T-cell targets in the spike protein suggest that pentapeptides – rather than hexapeptides – might be the targets of T-cell recognition (Cohen’s d 20.50 vs. 16.20). In contrast, the opposite was suggested by the analyses of overrepresentation of potential targets of recognition which are present in peptides that elicited T-cell response within all potential targets of T-cell recognition (Cohen’s d 1.02 vs. 1.97). Earlier proteomic analyses indicated that pentapeptides, and not hexapeptides, are the primary targets of the T-cells recognition. For example, a comparison of sizes of peptide vocabularies of 38 parasitic and 33 nonparasitic species showed that parasites of vertebrate hosts have impoverished pentapeptide vocabulary but enriched hexapeptide vocabulary (5). Similarly, the relative genetic distance between the horseshoe bat and SARS-CoV-2 vocabularies and between human and SARS-CoV-2 spike protein vocabulary was smaller for pentapeptides than for hexapeptides (7).

These discrepancies between previous and current results could be explained as follows: pentapeptides are recognized by T-cell receptors, but longer peptides are needed for attachment in the MHC protein groove (10). The specificity of a bond between MHC protein and the bordering parts of pentapeptides recognized by T cells might be relatively low; it may be so that several (though definitely not all) amino acids can border the focal pentapeptide to allow its attachment to the particular allele of the MHC gene. If a clone of T cell recognizes a particular pentapeptide, then probably several peptides containing different hexapeptides which include this pentapeptide are also recognized as immunogenic in immunological assays. It is possible that due to these ‘pseudoreplications’, statistical and Monte Carlo tests more easily detect enrichment of immunogenic peptides by hexapeptide than by pentapeptide targets. Due to this effect, we would more easily detect hexapeptides as targets of T-cell recognition even if pentapeptides were the genuine targets of T-cell recognition.

Regardless of whether pentapeptides or hexapeptides are being recognized by the T cells, the impoverishment of peptide vocabularies can be more easily recognized on the level of pentapeptides than on the level of hexapeptides. As discussed earlier (5), there are at least two processes that most likely affect the size of hexapeptide vocabularies. The first is the elimination of hexapeptides. This process necessarily accompanies the elimination of pentapeptides, because elimination of a pentapeptide can theoretically result in the elimination of as many as 40 different hexapeptides containing that particular pentapeptide. As argued earlier, the rate of elimination of hexapeptides from proteins during parasite’s adaptation to a new host is probably faster than the rate of elimination of pentapeptides (7). Another process, which runs in the opposite direction, is the enrichment of hexapeptide vocabulary whose aim is to compensate for the shortage in the number of pentapeptides. To build up biologically active proteins using a smaller collection of (pentapeptide) building blocks, the parasites must use these blocks more inventively – and this automatically results in a richer hexapeptide vocabulary.

The fact that many potential T-cell targets were not detected in any immunogenic peptides can be explained easily, as discussed above. More interesting is the existence of about one-half of immunogenic peptides which do not contain any potential pentapeptide T-cell targets. As discussed in the previous paragraphs, it is possible that hexapeptides, not pentapeptides – or, even more probably, both hexapeptides and pentapeptides – are the genuine targets of T-cell recognition. It can be argued that experimental approaches (such as crystallography, site-directed mutagenesis, and binding inhibition experiments), rather than bioinformatics, can tell us whether pentapeptides, hexapeptides, or both are recognized by T cells. Such experimental studies have already identified the amino acids which are in physical contact with the hypervariable CDR3 loop of T-cell receptors (TCR), as well as those which are necessary for the binding or correct positioning of a peptide in the groove of MHC molecules for a large number MHC-peptide-TCR complexes (11). Even these direct methods, however, are not omnipotent. Not all identified amino acids are necessarily part of the hexapeptide or pentapeptide that allowed T-cell recognition. To repeat: the presence of a parasite’s peptide that is absent in the host vocabulary is a necessary but not sufficient condition of T-cell recognition.

Another possible explanation for the total absence of predicted targets in some immunogenic peptides is that human T cells recognize even some pentapeptides of human origin as non-self. Elimination of potentially autoreactive T cells in the process of negative selection during their maturation does not work with a hundred percent efficiency. If some pentapeptides are present in only weakly expressed human proteins, or in proteins weakly expressed in the cells of the human thymus, then the corresponding T cells might survive the passage through the thymus. Such peptides then might become the targets of T-cell response when they are present in strongly-enough expressed viral proteins. This mechanism can play a role in some types of autoimmune disorders, because infection by a parasite can result in an expansion of initially rare potentially autoreactive T-cell clones (12) and the existence of certain classes of regulatory T cells might be part of adaptation of the immune system to such situations.

It is also probable that, due to polymorphism in most human proteins, some pentapeptides are missing in the protein vocabulary of some individuals, including the individual whose proteome was used for the construction of peptide vocabularies in the present study (F _000001405.39). In future studies, or when designing vaccines, it might be helpful to prepare a more representative human peptide vocabulary by using the proteomes of several individuals of different ethnic origin. In fact, one could even construct different vocabularies used for identification of potential T-cell recognition targets for people of different geographic or ethnic origin. Such approaches could mitigate the effect of genetic polymorphism in human protein-coding genes.

An efficient vaccine must contain not only the antigen that could bind to B-cell receptors (to membrane-bound and soluble immunoglobulins) but also peptides that could bind to receptors of helper T cells. Current algorithms for predicting which protein will elicit a T-cell response are based on rules derived from empirical studies (13–15). These algorithms provide useful outputs in searches for peptides which stimulate cytotoxic T-cell response but perform less well in searches for peptides which stimulate helper T-cell response (8). The efficiency of these algorithms is somewhat questionable when it comes to detection of cytotoxic T-cell targets. For example, in the study by Tarke *et al*., only 454 (8%) of all 5,600 predicted epitopes induced cytotoxic T-cell response. For comparison: from the 983 pentapeptides identified as potentially immunogenic by our method, 30% were present in peptides empirically found to elicit T-cells response. For those originating from the spike protein, as much as 70% were present in peptides that elicited the T-cell response.

New method of predicting potentially immunogenic peptides is based on rules derived from a theory, i.e., it is based on our knowledge of the universal mechanism of discrimination between self and non-self by vertebrate adaptive immune system. We recommend using our theory-based method when efficient empirically based algorithms are unavailable, for instance, during the development of veterinary vaccines. In all other cases, we recommend using it in combination with empirically based methods.

## Online Methods

The peptide vocabularies of human and SARS-CoV-2 were prepared as described earlier (7). The list of penta-, hexa-, and heptapeptides which were present in SARS-CoV-2 proteins but absent in human proteins, i.e., the list of potential targets of T-cells recognition, was assembled using the ImunDist 2.0 program [https://figshare.com/articles/software/ImunDist/17711474]. Tarke *et al*. had published lists of T-cell response targets in supplementary tables S5 and S8 of their paper (8). A subset of these genuine targets of T-cells response, which contained some of our potential targets of T-cells recognition, was obtained by the mcSortStrings 1.0 program, while the subset of potential targets of T-cell recognition contained (as a substring) in at least one genuine target of immune response was calculated using the mcPeptides 1.0 program [10.6084/m9.figshare.20981719]. These two programs (R scripts) were also used to perform one-sided Monte-Carlo tests (see Fig. 3). All lists were generated for the SARS-CoV-2 proteome as a whole and then also separately for the spike protein and ‘the rest of the proteome’, i.e., for the ten other SARS-CoV-2 proteins screened for T-cell targets in the (8) study (products of genes M, N, nsp3, nsp4, nsp6, nsp12, nsp13, nsp16, ORF3a, ORF8).

In both Monte Carlo tests, we have 10,000 times randomly selected a set of pseudo-targets from all overlapping peptides of a given length that were present in the SARS-CoV-2 proteome; these random sets were always of the same size of the actual set of potential targets of T-cell recognition detected earlier by the ImunDist 2.0 program. In the first Monte Carlo test (performed with mcSortStrings), we computed significance as the fraction of 10,000 random sets containing peptides which were present in a higher or equal number of genuine targets of T-cell response than peptides of the actual set of potential targets of T-cell recognition. In the second Monte Carlo test (performed with mcPeptides), we computed significance as the fraction of 10,000 random sets with a higher or equal number of peptides contained in the genuine targets of T-cell response than the peptides contained in the genuine set of potential targets of T-cell recognition. No correction for multiple (two) tests was done during post hoc testing (investigation of whether helper or cytotoxic T-cells were responsible for the observed effect).

## References

1. Lanzavecchia A. Antigen-specific interaction between T-cells and B-cells. Nature. 1985;314:537–9.

2. Jagger A, Shimojima Y, Goronzy JJ, Weyand CM. Regulatory T cells and the immune aging process: A mini-review. Gerontology. 2014;60(2):130–7.

3. Foulsham W, Marmalidou A, Amouzegar A, Coco G, Chen YH, Dana R. Review: The function of regulatory T cells at the ocular surface. Ocul Surf. 2017;15(4):652–9.

4. Najafi M, Farhood B, Mortezaee K. Contribution of regulatory T cells to cancer: A review. J Cell Physiol. 2019;234(6):7983–93.

5. Zemkova M, Zahradnik D, Mokrejs M, Flegr J. Parasitism as the main factor shaping peptide vocabularies in current organisms. Parasitology. 2017;144:1–9.

6. Flegr J. Pozor, Toxo! Tajná učebnice praktické metodologie vĕdy (Watch out for Toxo! The secret guide to practical science). Prague: Academia; 2011.

7. Flegr J, Zahradník D, Zemková M. Thus spoke peptides: SARS-CoV-2 spike gene evolved in humans and then shortly in rats while the rest of its genome in horseshoe bats and then in treeshrews. Communicative & Integrative Biology. 2022;15(1):96–104.

8. Tarke A, Sidney J, Kidd CK, Dan JM, Ramirez SI, Yu ED, et al. Comprehensive analysis of T cell immunodominance and immunoprevalence of SARS-CoV-2 epitopes in COVID-19 cases. Cell Rep Med. 2021;2(2).

9. Cohen J. Statistical power analysis for the behavioral sciences. New York, New York 10003: Academic Press Inc.; 1977.

10. Vyas JM, Van der Veen AG, Ploegh HL. The known unknowns of antigen processing and presentation. Nature Reviews Immunology. 2008;8(8):607–18.

11. Garcia KC, Adams EJ. How the T cell receptor sees antigen--a structural view. Cell. 2005;122(3):333–6.

12. Cole DK, Bulek AM, Dolton G, Schauenberg AJ, Szomolay B, Rittase W, et al. Hotspot autoimmune T cell receptor binding underlies pathogen and insulin peptide cross-reactivity. J Clin Invest. 2016;126(9):3626.

13. Jurtz V, Paul S, Andreatta M, Marcatili P, Peters B, Nielsen M. NetMHCpan-4.0: Improved Peptide-MHC Class I Interaction predictions integrating eluted ligand and peptide binding affinity data. J Immunol. 2017;199(9):3360–8.

14. Karosiene E, Rasmussen M, Blicher T, Lund O, Buus S, Nielsen M. NetMHCIIpan-3.0, a common pan-specific MHC class II prediction method including all three human MHC class II isotypes, HLA-DR, HLA-DP and HLA-DQ. Immunogenetics. 2013;65(10):711–24.

15. Paul S, Arlehamn CSL, Scriba TJ, Dillon MBC, Oseroff C, Hinz D, et al. Development and validation of a broad scheme for prediction of HLA class II restricted T cell epitopes. J Immunol Methods. 2015;422:28–34.

